# Fronto-parietal networks shape human conscious report through attention gain and reorienting

**DOI:** 10.1101/2022.04.10.487690

**Authors:** Jianghao Liu, Dimitri J. Bayle, Alfredo Spagna, Jacobo D. Sitt, Alexia Bourgeois, Katia Lehongre, Sara Fernandez-Vidal, Claude Adam, Virginie Lambrecq, Vincent Navarro, Tal Seidel Malkinson, Paolo Bartolomeo

**Affiliations:** Sorbonne Université, Inserm, CNRS, Paris Brain Institute, ICM, Hôpital de la Pitié-Salpêtrière, 75013 Paris, France; Dassault Systèmes, Vélizy-Villacoublay, France; Licae Lab, Université Paris Nanterre, Nanterre, France; Department of Psychology, Columbia University in the City of New York, NY, 10027, USA; Laboratory of Cognitive Neurorehabilitation, Faculty of Medicine, University of Geneva; 1206 Geneva, Switzerland; CENIR - Centre de Neuro-Imagerie de Recherche, Paris Brain Institute, ICM, Hôpital de la Pitié-Salpêtrière; 75013 Paris, France; AP-HP, Pitié-Salpêtrière Hospital, Epilepsy Unit, 75013, Paris, France; AP-HP, Pitié-Salpêtrière Hospital, Clinical Neurophysiology Department, 75013, Paris, France

## Abstract

How do attention and consciousness interact in the human brain? Rival theories of consciousness disagree on the role of fronto-parietal attentional networks in conscious perception. We recorded neural activity from 727 intracerebral contacts in 13 epileptic patients, while they detected near-threshold targets preceded by attentional cues. Unsupervised clustering revealed three patterns: (1) Attention-enhanced conscious report accompanied sustained right-hemisphere fronto-temporal activity, in networks connected by the superior longitudinal fasciculus (SLF) II-III, and late accumulation in bilateral dorso-prefrontal and right-hemisphere orbitofrontal cortex (SLF I-III). (2) Attentional reorienting affected conscious report through early, sustained activity in a right-hemisphere network (SLF III). (3) Conscious report accompanied left-hemisphere dorsolateral-prefrontal activity. Task modeling with recurrent neural networks identified specific excitatory and inhibitory interactions between attention and consciousness, and their causal contribution to conscious perception of near-threshold targets. Thus, distinct, hemisphere-asymmetric fronto-parietal networks support attentional gain and reorienting in shaping human conscious experience.

**One-Sentence Summary:** Intracerebral recordings, tractography and modeling reveal the interaction of attention and consciousness in the human brain.

## Introduction

How does attention impact consciousness? Despite decades of research^1–4^ the interactions between these two core cognitive concepts remain unclear. Rival theories of consciousness vary in the role ascribed to attention in conscious perception, both conceptually and neurally, and especially on the role of fronto-parietal (FP) networks^5^, which are strongly associated with attention processing^6, 7^. Consequently, the proposed relationship between attention and consciousness is one of the key distinctions between consciousness theories^8^. Some theories explicitly include attention as a modulating factor for consciousness^4, 9, 10^. According to the global neuronal workspace hypothesis (GNWH), to attain conscious processing near-threshold stimuli must be attended and consequently receive top-down amplification^10^. In conscious perception, neural information is sustained and globally broadcasted across the brain, with an important role for dorsolateral prefrontal cortex (PFC) and inferior parietal cortex^11^. This idea about the relationship between attention and consciousness is also consistent with the gateway hypothesis^9^. In other theories, consciousness depends partly or not at all on attention-associated regions in the frontal or parietal cortex, without an explicit conceptual role for attention. For example, the integrated information theory^12^ postulates that conscious information is integrated in a temporo-parietal-occipital “hot zone”^13^, and the recurrent processing theory^14^ holds that conscious experience emerges from reverberating activity in sensory areas. Both theories postulate that FP networks contribute to post-conscious cognitive processing and task relevance of targets, such as motor planning or verbal report^15, 16^, and claim that attention and consciousness are distinct both conceptually and neurally. Alternatively, attention and consciousness could be implemented by distinct neural mechanisms but have cumulative influence on the behavioral report. The cumulative influence hypothesis postulates the existence of an interaction between attention and consciousness solely at the behavioral level, but not in neural activity^4^.

One reason for this theoretical divergence might be that attention is a heterogeneous group of neurocognitive processes^17, 18^. For example, endogenous and exogenous spatial attention refer to distinct behavioral and neural dynamics in partially overlapping neural substrates^19, 20^. The available evidence suggests that endogenous, or top-down, attention has little role in supporting conscious perception^1, 4, 21, 22^, while exogenous, or stimulus-driven, attention is a necessary, although not sufficient, condition for conscious perception^23, 24^. In neurotypical participants, exogenous cues near the spatial location of an upcoming near-threshold target increase the target’s conscious detection^22, 23, 25^. This increase is accompanied by a higher activation of the dorsal FP attentional network^26^ for seen compared to unseen targets at attended locations^27^. Moreover, neurological patients with signs of spatial neglect^28^ display a systematic pattern of association between right-biased exogenous attention and unawareness of left-sided events^29^. This clinical evidence strongly suggests a specific role for right hemisphere FP attention networks in conscious processing.

Despite this converging evidence, which points to the modulation of consciousness by exogenous attention in both behavior and neural activity, it is still unclear where this interaction occurs in the brain and how different brain networks interact to achieve this effect. Further, the spatiotemporal resolution of neuroimaging techniques like fMRI and EEG used so far to study these questions is too rough for establishing the neural basis of the rapid and dynamic exogenous attention modulation of conscious perception. Facing the divergence of theoretical predictions, and the resolution limitations of the evidence collected hitherto, we decided to use a data-driven approach to try to establish the dynamics of the neural interactions between attention and consciousness on a fine scale, by taking advantage of the excellent spatiotemporal resolution of human intracerebral EEG (iEEG) recordings. We tested empirically using unsupervised clustering the division of functional clusters of iEEG contact based on the neural temporal patterns of the interaction between exogenous attention and conscious report, experimentally manipulated in a detection task. This allowed us to reveal the brain areas supporting different patterns of attention/consciousness interaction. We further employed white-matter tractography to collect connectomic evidence on the network architecture of the functional clusters. Finally, we used recurrent neural network models to casually examine the computations in the neural clusters and elucidate inter-cluster interactions, which critically contribute to behavior.

## Results

### Behavioral results: Cue validity modulates target detection

We recorded neural activity from 727 intracerebral contacts in 13 patients receiving presurgical evaluation of drug-resistant epilepsy (age 34.7 ± 8.7 years; 7 women). Patients performed a near-threshold target detection task^30^ (Fig. 1A), in which they attempted to detect left- or right-tilted, near-threshold Gabor patches (the targets), presented either left or right of fixation. The target was preceded by supra-threshold peripheral non predictive visual cues, which appeared either on the same side as the subsequent target (Valid cues) or on the opposite side (Invalid cues). All conditions (target side, cue validity) were randomly interleaved, with 20% of cue-only, “catch” trials, where no target was presented. Individual Gabor contrasts based on an individual calibration procedure were used across all conditions. Participants had to discriminate the direction of the Gabor’s tilting, and subsequently report the presence or absence of the Gabors. They were informed that cues did not predict the location of the upcoming targets.

**Fig 1.**
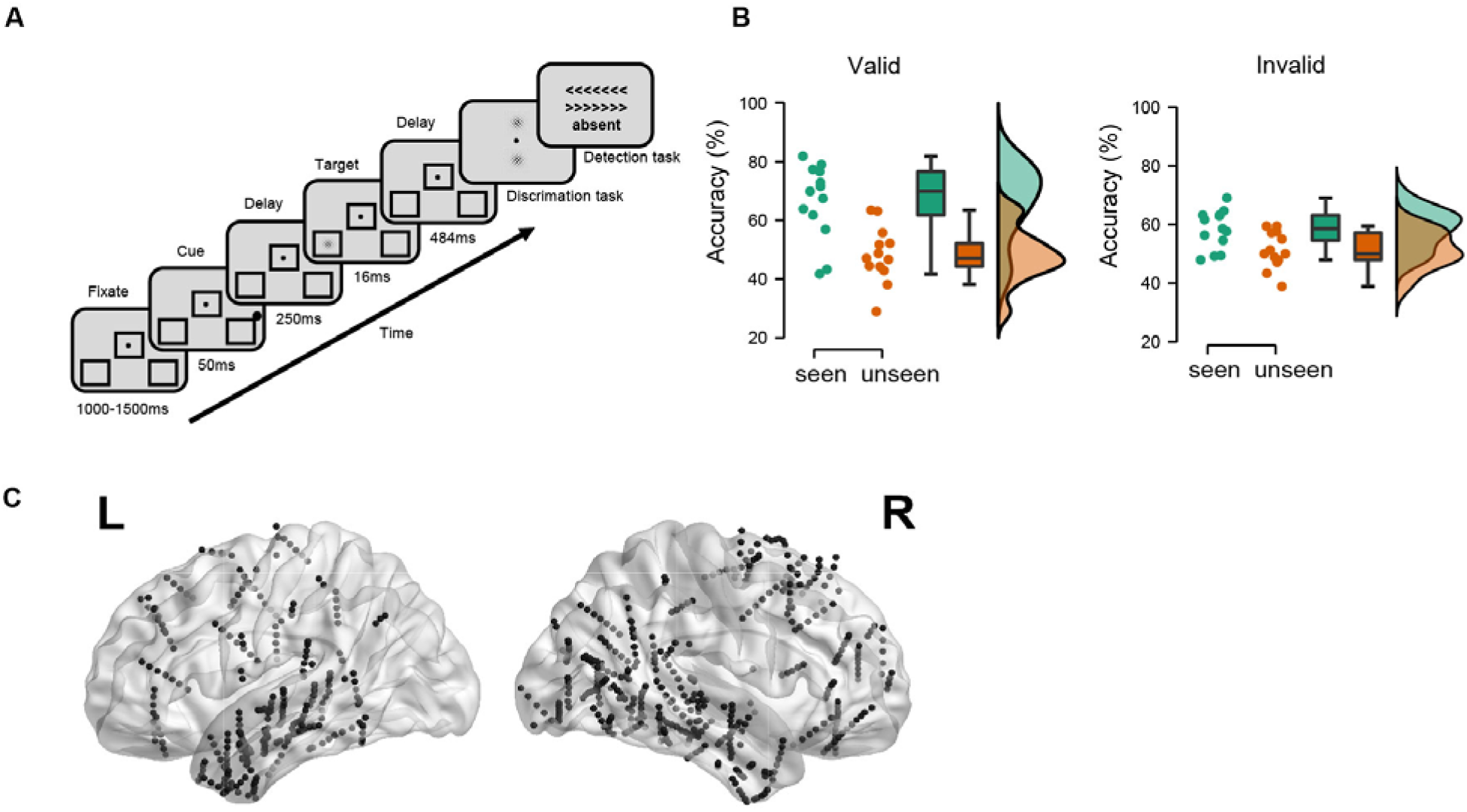
Near-threshold target detection task and human intracerebral recordings. **A.** After a fixation dot lasting 1,000-1,500ms, a peripheral non-predictive dot cue occurred for 50ms, followed by a left-sided or right-sided near-threshold tilted Gabor target presented for 16ms. After a delay of 484ms, participants discriminated the direction of tilting and reported the presence or absence of the Gabor. In 20% of trials (“catch” trials), the Gabor target was absent. Individual contrasts based on an individual calibration procedure were used across all conditions. **B.** Target discrimination accuracy. Dots represent individual performance. Chance level is at 50%. **C.** The 727 intracerebral contacts pooled from 13 epileptic patients in MNI space, see detailed localization label in Table S1.

To see whether exogenous attention modulates the conscious perception of targets in the discrimination task, we performed a two-way ANOVA with the factors of cue validity and conscious report on the percentage of accurate responses. As expected, participants were more accurate in discriminating target direction when the targets were reported to be perceived than when they went unseen (main conscious report effect: 𝑭_1,24_ = 24.74, *p* < 0.001, η² = 0.315). Although the main validity effect did not reach significance (𝑭_1,24_ = 1.04, *p* = 0.31, η² = 0.013), validity interacted with conscious detection, because validly cued targets were more likely to be reported than invalidly cued ones (𝑭_1,24_ = 4.77, *p* < 0.05, η² = 0.061, see Fig. 1B; post hoc *t*-test: valid vs. invalid for seen targets, t-value = 3.04, *p* < 0.05, Cohen’s d = 0.84). Signal detection theory (SDT) analysis^31^ showed that valid cues induced a higher detection rate (pairwise *t*-test, *p* < 0.05, Cohen’s d = 0.58), a higher false alarm rate (pairwise *t*-test, *p* < 0.01, Cohen’s d = 0.87), and a more liberal response criterion (pairwise *t*-test, *p* < 0.01, Cohen’s d = 1.03) than invalid cues (Fig. S1A). However, no significant difference emerged in sensitivity (pairwise *t*-test, *p* = 0.52). No significant interaction effects were found in response times.

### Five neural clusters associated with consciously perceived targets form three patterns of interaction with attention

After discarding epileptic artifacts, there were 727 usable contacts with bipolar montage pooled across all participants (288 in the left hemisphere; 439 in the right hemisphere; Fig. 1C, see Table S1 for details). For each contact, we extracted high-frequency broadband power (HFBB; 70-140 Hz), which is generally considered as a proxy of neuronal population activity^32, 33^, but also see Ref.^34^. We then computed target-locked mean normalized HFBB across the eight experimental conditions (2×2×2 design: target side [ipsilateral/contralateral] x cue validity [valid/invalid] x conscious report [seen/unseen]). Neural activity components of all contacts were visualized in a two-dimensional t-distributed stochastic neighbor embedding (t-SNE) (Fig. S1B).

Next, we applied a trajectory *k*-means clustering method^35^ to identify the main groups of contacts that carry cue validity and conscious report information (see Methods). This procedure allowed us to group intracerebral contacts based on their temporal profile of neural activities across the experimental conditions. For each contact, we computed the temporal trajectory in the eight-dimensional condition space, i.e. the path of each contact’s HFBB power over time across all conditions (target side, cue validity, conscious report). Using *k*-means clustering, each trajectory was then assigned to the cluster with the nearest trajectory-centroid, by iteratively minimizing within-cluster Manhattan distances. A ten clusters solution reflected the highest average silhouette score serving as the evaluation criterion of the clustering results (Fig. S2A). We explored how our experimental manipulation of attention and consciousness influenced the clusters’ activity. For each cluster, we performed time-resolved 3-way ANOVAs in both the cue-target period (from −300ms to target onset) and the post-target period (from target onset to 500ms post-target) with the factors of target side, validity, and conscious report. Five of the ten clusters showed a main effect of conscious report, with higher levels of activity for seen than for unseen targets (all *ps* < 0.018). The number of contacts in these clusters was stable (Fig. S2D) and their cluster-level temporal profiles were similar across *k*-means solutions with varying numbers of clusters. We thus focused on these clusters for further analyses (see Table S1 for details about anatomical localization in each cluster). The remaining five clusters didn’t show any significant effects and were not included in further analysis. We then explored how exogenous attention affects conscious reports of the targets in the five clusters, by examining the interaction between Cue validity and Conscious report (valid/invalid x seen/unseen) in the above mentioned time-resolved ANOVA. This analysis revealed 3 distinct patterns of neural activity. We will describe the interaction patterns as well as the main effect of conscious report in the cue-target and post-target periods for the five clusters (Fig. 2A).

1. The first interaction pattern encompassed three out of the five clusters and showed enhanced conscious report effect for validly cued targets compared to invalidly cued targets. The first cluster (hereafter: the Visual cluster, 42 contacts) showed an early transient post-target effect, with stronger activation for seen targets compared to unseen ones (90-350ms and 380-430ms, all *ps* < 0.003). Additionally, the Visual cluster showed an interaction between target side and cue validity in the cue-target period (60-210ms after cue onset, all *ps* < 0.006, Fig. S2E), with higher neural activity for contralateral than for ipsilateral cues. However, there was no significant Target side effect in the post-target period, perhaps because of the low, near-threshold intensity of the targets. No significant three-way interaction emerged (all Fs < 8.17, *ps* > 0.20). This cluster mainly consisted of contacts in the right posterior temporoparietal areas (there was no available electrode in the homolog areas of the left hemisphere). The second cluster (Sustained cluster, 148 contacts) showed an effect of consciousness both in early cue-elicited (−140 to −90ms before target onset, all *ps* < 0.016) and in later, target-related sustained neural activity (160-200ms, 240-300ms, 340-430ms, and 450-500ms post target, all *ps* < 0.003). The contacts of the Sustained cluster were mainly located in the bilateral temporal cortex, the right angular gyrus (AG), and the right PFC, around the central portion of the right superior frontal gyrus. The activity of the third cluster (Late accumulation cluster, 67 contacts) increased over time in the post-target period for seen targets (300-500ms, all *ps* < 0.005), but there was no significant conscious report effect in the cue-target period. Most of the contacts in the Late accumulation cluster were located in the bilateral PFC, around the left inferior frontal gyrus (IFG), the right orbitofrontal cortex (OFC) and the caudal portion of the right superior frontal gyrus. Similar to the attentional enhancement on conscious report in behavior, there was an attention-related enhancement interaction with conscious report in three neural clusters (Visual cluster: 190-220ms, all *ps* < 0.03; Sustained cluster: 270-330ms, all *ps* < 0.002; Late accumulation cluster: 360-430ms, all *ps* < 0.006). The amplitude of this interaction did not significantly differ across the three clusters by direct comparison (three-way ANOVA with the factors of cluster x validity x conscious report, all *ps* > 0.30), possibly due to faint target contrasts. A further explorative analysis compared this enhancement, by Cohen’s d values which were derived from time-resolved t-test for the interaction contrast (seen valid - unseen valid) - (seen invalid - unseen invalid), around the time points where the interaction was significant. The result showed an increasing effect size gradient from the Visual cluster to the Sustained and Late accumulation clusters (Fig. S3D, one-way ANOVA: 𝑭_1,24_= 3.83, *ps* < 0.05, η² = 0.15, linear polynomial contrast, *p* < 0.05, t = 2.48), suggesting an increasing attention modulation on conscious report along these clusters.
2. A Reorienting cluster (19 contacts) showed a reversed pattern, with higher activity for seen targets after invalid cues than for seen targets after valid cues, early after target onset (160-190ms, all *ps* < 0.002). This cluster presented fast activity in the cue-target period (−250 to −220ms and −130 to −90ms before target onset, all *ps* < 0.015), as well as early in the post-target period (170-230ms, all *ps* < 0.001), that resulted in “seen” reports for uncued targets. This cluster was localized in the right hemisphere temporoparietal junction (TPJ) / IFG. Activity in this cluster was sustained after target onset, then showed a transient peak around 180ms only for invalid seen targets; however, for seen targets after valid cues, activity decreased once targets appeared. This cluster thus was likely to reflect reorienting of attention from the invalid cue to the opposite target ^26^.
3. A Conscious report cluster (38 contacts) showed a late sustained neural activity selective for seen targets (310-450ms, all *ps* < 0.018), independent of cue validity. This cluster contains contacts from the left posterior portion of dorsolateral PFC, around the left frontal eye field, and the bilateral posterior temporal area.

**Fig 2.**
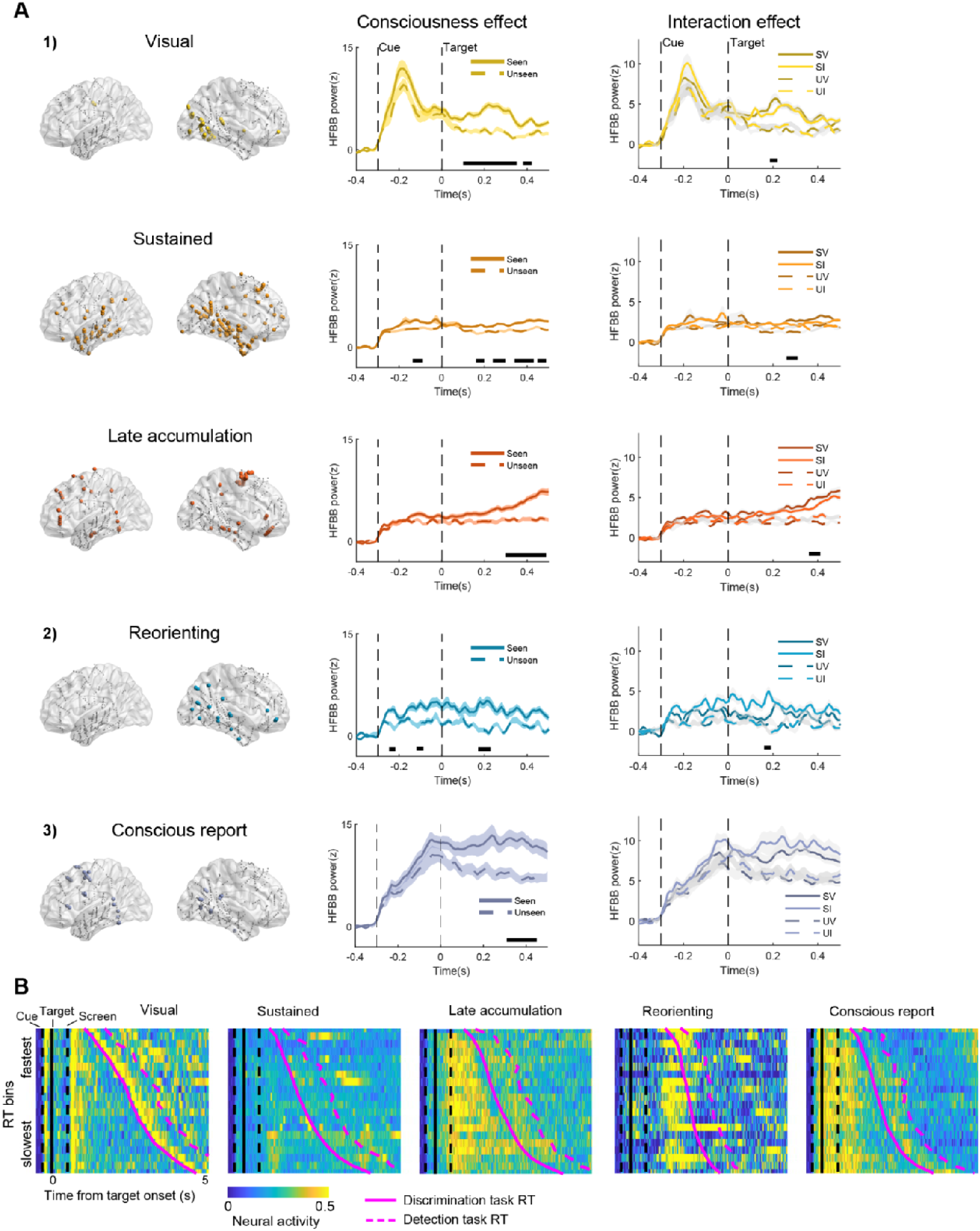
Trajectory k-means clustering revealed five clusters showing a conscious report effect and three patterns of interaction of exogenous attention with conscious report. **A.** Three neural patterns of the validity X conscious report interaction: 1) three clusters (Visual, Sustained, Late accumulation) showed enhanced conscious report effect for the validly cued targets. Left panels, cluster contact localization; middle panels, comparison of target-locked neural activity for seen and unseen trials in each cluster, black horizontal bars underneath the traces indicate *p* < 0.05, Holm-Bonferroni corrected, shading is ± SEM across electrodes; right panels, comparison of interaction effect, SV: seen valid; SI: seen invalid; UV: unseen valid; UI: unseen invalid, black horizontal bar for all *p*s < 0.05, Holm-Bonferroni corrected, gray shading is ± SEM across electrodes. 2) Reorienting cluster showed early sustained neural activity for invalidly cued targets. 3) a late Conscious report cluster differentiated seen from unseen targets independent of attention. Note that the Sustained cluster and the Reorienting cluster also showed significant conscious target report effects during the cueing period, before target occurrence. **B.** Visual- and RT-modulation of target-locked neural activity. RT bins were sorted according to their mean RT from fastest to slowest, with neural activity pooled across contacts in each cluster. Magenta full curve shows mean discrimination task RT, followed by dashed magenta curve for mean detection task RT. Neural activity in the Visual cluster synchronized with visual stimuli (cue, response screen and screen switching after discrimination task). Late accumulation cluster showed sustained neural activity until the response. Conscious report cluster exhibited sustained neural activity locked to the visual percept, but not to the report.

The differences of temporal components of the three neural patterns in the interaction between exogenous attention and consciousness can be visualized in a two-dimensional t-distributed stochastic neighbor embedding (t-SNE) decomposition (see Fig. S3A). This visualization corroborates the separation of neural dynamics of the five clusters in the interaction between attention and conscious report.

We then sought to understand the functional roles of the clusters by relating the neural activities to the behavioral responses. In each cluster, we divided the trials (pooled across conditions) into 20 quantiles according to their response time (RT) in the discrimination task (see Methods; note that in the discrimination task patients had to wait for the onset of a response screen in order to respond). We tested the relation of the neural activities in RT-bins using time-resolved one-way repeated measures ANOVA, in a time window from target onset to 1,000ms, to avoid the influence of neural activity in the subsequent trial. The Late accumulation cluster showed target-locked sustained neural activity in trials evoking slower RTs (340-1,000ms, all *ps* < 0.003; see Fig.S3B), but not in trials with faster RTs. The other clusters had no or only transient (less than 300ms) RT effects for targets. We then compared the neural activity of the 10 fastest RT bins with those of the 10 slowest RT bins. The Late accumulation cluster showed greater accumulating and sustained activity for slower RTs responses than faster ones (Fig. S3C). Interestingly, the Visual cluster showed a stimulus-locked higher neural activity for faster RTs, perhaps because faster responses resulted from stronger evidence for target presence. No significant effect emerged in other clusters. Next, we visualized the neural activity across RT bins over a longer time window, which contains the neural information until the button press (target onset to 5,000ms, see Fig. 3B). Here, activity might be affected by the subsequent trial; hence, we report only observational findings without statistical testing. We observed that the transient response elicited in the Visual cluster was locked to visual modulations (i.e., the appearance of cue, target, screen, and switching of the display after the discrimination task). The Late accumulation cluster showed accumulated and sustained activity until report. The Reorienting cluster shows a late transient response. The Conscious report cluster elicited sustained neural activity, which was locked to the visual percept and was not associated with the time of report. To test whether these activities were related to motor preparation, we computed the beta band power (16 - 28 Hz), which typically decrease with motor planning^36^. There was no sign of decreasing beta activity in the Late accumulation cluster and in the Conscious report cluster. This result suggests that motor preparation had little role in eliciting these clusters (Fig. S3E).

**Fig 3.**
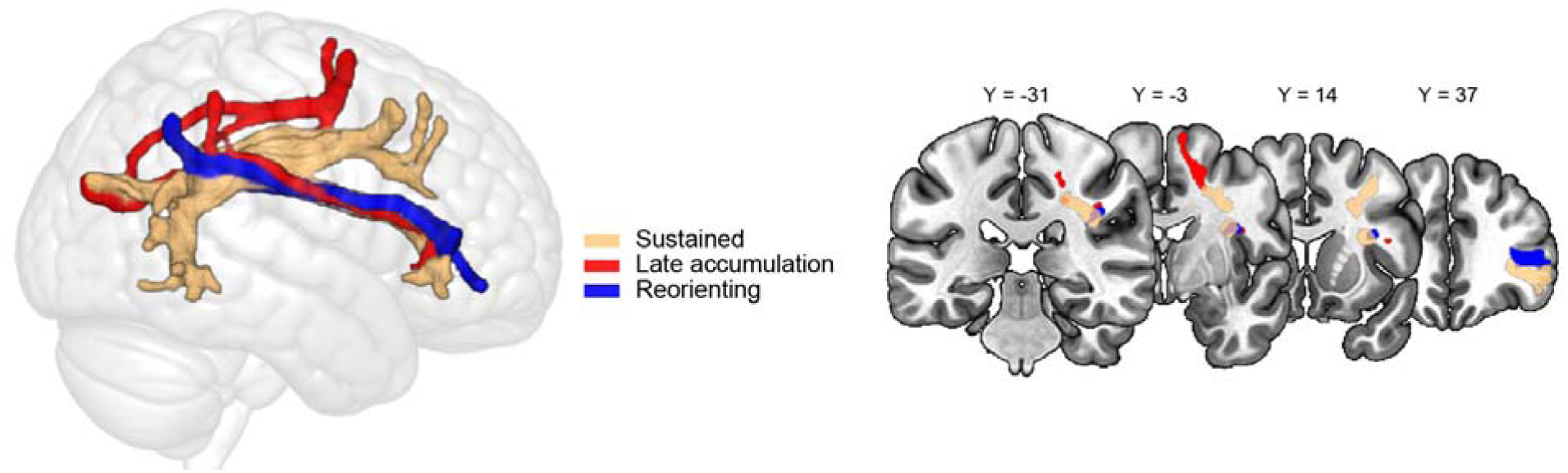
FP white matter tracts connecting contacts in the right hemisphere. Tractography t-maps, showing the significant white matter voxels (threshold-free cluster enhancement-based non-parametric *t*-test, *p* < 0.05), which connect frontal and parietal contacts within each cluster

### Fronto-Parietal white matter tracts connecting contacts in the right hemisphere

To specify the anatomical connections between contacts within each cluster, we performed white matter tractography analysis paired with probability maps in 176 healthy individuals from the Human Connectome database^37^. Fig. 3 displays the white matter tracts connecting frontal contacts and parietal contacts within each cluster. This result suggests that our unsupervised cluster analysis mapped on existing anatomo-functional networks^7^. We examined the frontal association tracts described in Ref.^38^, and found that our frontal and parietal contacts were connected by branches of the superior longitudinal fasciculus (threshold-free cluster enhancement-based non-parametric t-test, *p* < 0.05). Specifically, the Sustained cluster was mainly connected by the right superior longitudinal fasciculus (SLF) II (43.4%) and III (36.8%). The Late accumulation cluster was mainly linked by the right SLF I (39.4%) and III (25.1%). The Reorienting cluster was connected by right SLF III (84.2%). No statistically significant tracts emerged from the analysis of the Visual and the Conscious report cluster.

### Task modeling with recurrent neural network

To better understand the relation between activity dynamics and behavior, we simulated the task with a recurrent neural network (RNN) model (Fig. 4A). We separately modeled left- and right-sided visual stimuli as two noisy signal inputs. The model had a single layer containing 50 units recurrently connected to one another, of which 80% were excitatory units and 20% were inhibitory units. The network was trained by back-propagation to produce two different outputs, one for each side, about the presence or absence of the target. Similarly to the human task, two RNN outputs were combined to measure a discrimination and a detection performance. The task conditions were also similar to the human task, and included valid trials, invalid trials, and 20% catch trials (see examples of stimuli inputs, hidden units dynamic and network outputs in Fig. 4B for valid trial and in Fig. S4A for invalid and catch trials).

**Fig 4.**
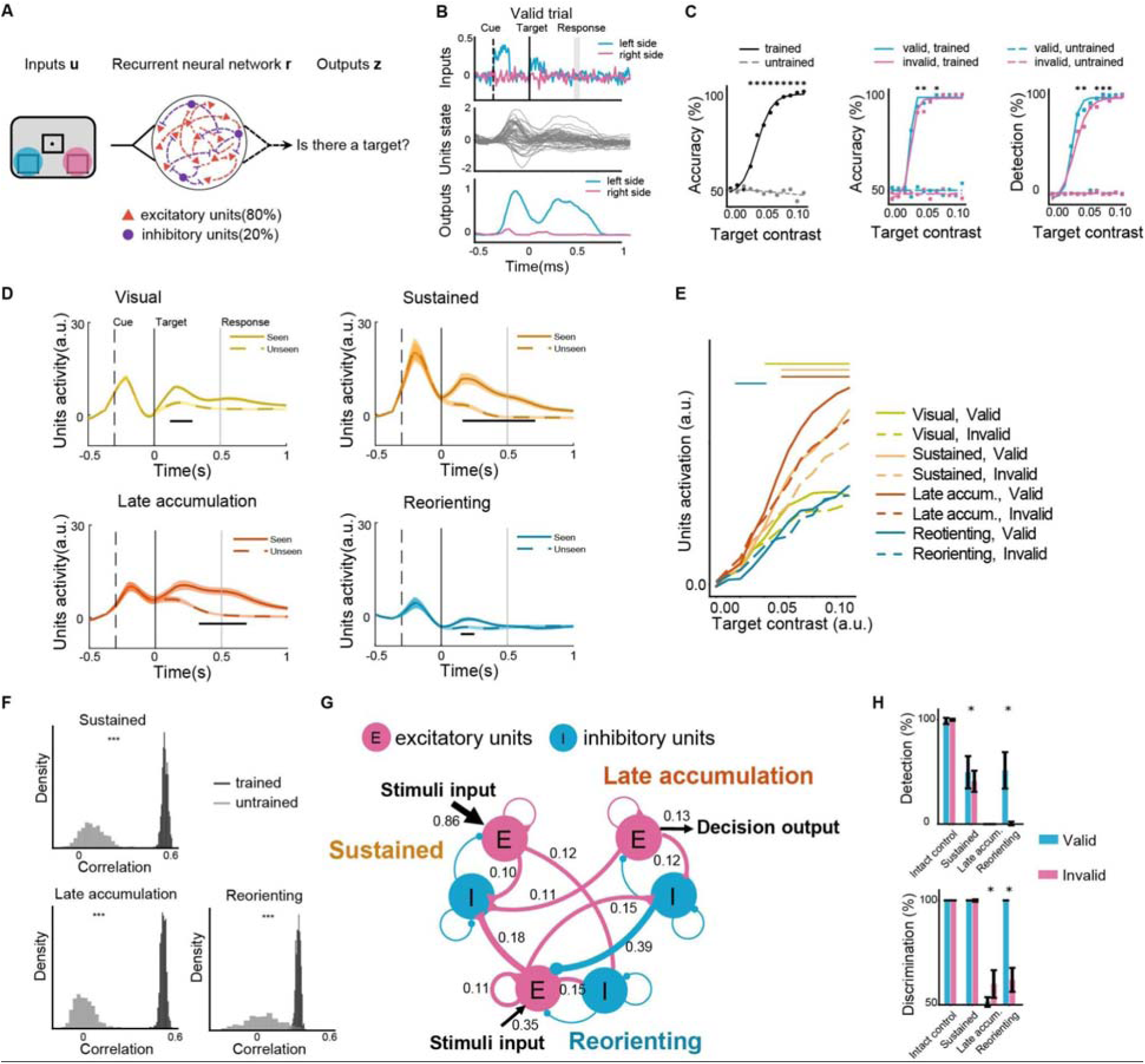
Task modeling with recurrent neural networks. **A.** Recurrent neural network model and task. Left- and right-sided visual stimuli were separately modeled as two noisy inputs. The model had a single layer containing 50 recurrent units. The network produced two outputs to decide whether a target was present on the left or on the right side. **B.** Example of a trial showing the task structure and hidden unit dynamic as well as outputs from a trained model. After a fixation period, cue signal was presented on either the left or the right side, followed by a target signal presented on either side or absent. In a valid trial, cue signal and target signal were presented on the same side. Gray vertical line indicates the response window. **C.** Task performance. Left panel: discrimination psychometric curve for all targets. Lines show the best-fitting logistic function to 12 target contrasts. Comparison of accuracy of the trained model to the untrained model by *t*-test, \**p* < 0.001, Bonferroni-corrected. Middle panel: discrimination psychometric curve. Comparison of discrimination accuracy for validly cued targets *versus* invalidly cued targets in trained model by *t*-test, \**p* < 0.05, Bonferroni-corrected. Right panel: detection psychometric curve for validly cued targets *versus* invalidly cued targets. **D.** Trajectory *k*-means clustering of model unit activities. Four unit clusters showed distinct temporal trajectories in seen *versus* unseen trials for intermediate target contrast level. Black lines for all \**p* < 0.05, Holm-Bonferroni corrected. a.u.: arbitrary unit. **E.** Comparison of unit activities for valid *versus* invalid trials in each cluster while increasing stimulus contrast. Late accum.: Late accumulation. Horizontal bar above the curves for all \**p* < 0.05, Holm-Bonferroni corrected. a.u.: arbitrary unit. **F.** Histograms of the correlation coefficient between the trajectory of neural activity and unit activity. Permutation tests, \*\*\**p* < 0.001 for trained model versus untrained model. **G**. Directed cluster connection graph for groups of excitatory and inhibitory units from three clusters. Magenta curves show excitatory connection and blue curves for inhibitory connection. Triangle or circle at the end of the connection line represents the destination. Numbers denote the weight of connection, with significance level controlled at *p* < 0.05 in one-sample *t*-test, Holm-Sidak corrected across groups of units. **H.** Effect of cluster’s lesions. Intact control: unlesioned model; Late accum.: Late accumulation; Reorienting. \**p* < 0.05 for the comparison of valid *versus* invalid trials by *t*-test.

The trained model displayed a detection psychometric curve, resembling typical human performance^39–41^. This curve differed from that characterizing an untrained model with random Gaussian connectivity weights (all *ps* < 0.001 for stimulus contrast above 0.03, see Fig. 4C, left panel). Further, in the trained model, the improvement of target discrimination and detection with valid cues emerged only at sufficiently high stimulus contrast levels (Fig. 4C, middle and right panels). Consistent with this task performance pattern, a t-SNE visualization of RNN units’ components showed a difference for valid *vs*. invalid trials only in intermediate or higher target contrasts, but not with lower contrast levels (Fig. S4B-C). Thus, all further analyses on RNN units were performed on the intermediate target contrast levels, corresponding to near-threshold targets in the human task.

To identify the temporal patterns of activity of the RNN hidden units, we used our trajectory *k*-means clustering method. Similar to the human results, the clustering analysis with silhouette evaluation resulted in five stable clusters (Fig. S4D) with different temporal trajectories. Four of these clusters showed stronger unit activity for seen targets than for unseen targets (Fig. 4D). Again similar to the human neural data, after an early, transient activity in a Visual cluster (8 units, 110-300ms, all *ps* < 0.017), there was sustained unit activity in a Sustained cluster (20 units, 140-750ms, all *ps* < 0.032), and late activity in a Late accumulation cluster (8 units, 300-730 ms, all *ps* < 0.038). Mirroring the model’s task performance, the higher the target contrast, the greater the activity in these clusters. Importantly, these clusters showed significantly enhanced activities for validly cued targets at sufficiently high stimulus contrast (Fig. 4E), akin to neural amplification. The fourth, Reorienting, cluster showed early transient activity related to “conscious” detection (11 units, 170-230ms, all *ps* < 0.019). A t-SNE visualization of unit activity of these clusters showed distinct unit component patterns for seen targets preceded by valid *vs*. invalid cues (Fig. S5).

To examine potential similarities between RNN clusters and human neural clusters, we used a temporal trajectory correlation analysis. RNN unit activity in the Sustained, Late accumulation, and Reorienting clusters in the trained model were significantly more similar to the equivalent human neural clusters than to the clusters obtained from the untrained model (Fig. 4F, one-sided permutation test, all *ps* < 0.001). We examined how these model clusters are interconnected, and the nature of their computation. In the trained model, we extracted input weights (sensory enhancement gain), connection weights between units, and output weights (report gain) (Fig. S4E). The model only constrained the total number of excitatory (E) and inhibitory (I) units. After RNN training, both excitatory and inhibitory units emerged in each cluster. After grouping separately excitatory and inhibitory units in each of the three clusters (a total of 6 groups), we computed a directed cluster connection graph by averaging unit-to-unit connection weights from one group (“pre-synaptic”) to each of the other groups (“post-synaptic”) and compared the resulting connections to random weights (Fig. 4G, one-sample *t*-test, all *ps* < 0.05, Holm-Bonferroni corrected). The stimuli input was mainly connected to the excitatory units in the Sustained cluster, which was associated with the conscious processing-Sustained intracerebral cluster. The Late accumulation excitatory units were connected to the decision output, confirming its role in decision making. Notably, the Reorienting excitatory units also received a branch of stimuli input and showed strong excitatory connection to the inhibitory units in the Sustained and in the Late accumulation clusters, reflecting the Reorienting cluster’s role in early target monitoring and in executing inhibitory control over stimuli processing and decision making units.

Finally, we lesioned each of the RNN clusters to assess their causal contribution to task performance. For each cluster, we set all the unit weights to zero, and monitored the change in task performance (Fig. 4H). Lesion of either the RNN Sustained cluster or the Late accumulation cluster decreased the percentage of detected targets (“report” units). Lesion of the Late accumulation cluster additionally impaired discrimination accuracy. Lesion of the Reorienting cluster led to a selective failure to detect and to discriminate invalidly cued targets (“reorienting” units). However, performance reverted to normal with very high target contrast levels, presumably because these contrast levels were sufficient to capture attention even without the contribution of the Reorienting cluster.

## Discussion

We combined human intracerebral EEG, white matter tractography, and computational modeling to elucidate the fine-scale spatiotemporal dynamics of brain networks underlying exogenous attention modulation of conscious report. Unsupervised temporal clustering revealed three patterns of neural activity fronto-parietal networks, diverging from the classical model of dorsal and ventral attentional networks^26^. The three neural patterns supported the interaction between exogenous orienting and conscious report: (1) a Sustained cluster showed attention-enhanced sustained activity for validly cued targets; (2) a Late accumulation cluster with progressively showed increasing activity until report; (3) a Reorienting cluster showed an early, sustained response to invalidly cued targets. Using RNN modeling, we discovered multiple clusters matching the identified neural clusters, clarifying the nature of inter-cluster interactions and uncovered the causal contribution of these clusters to behavior. Altogether, our behavioral, neural, and modeling findings consistently demonstrate that exogenous attention, the process that makes external salient stimuli stand out in a visual scene, modulates conscious report.

We demonstrated distinct neural dynamic patterns that implemented signal enhancement and attentional reorienting interacting with conscious report in three right-hemisphere FP networks connected by branches of SLF, as well as inter-network interactions.

At the behavioral level, valid cues increased both detection rate and discrimination accuracy of near-threshold peripheral target and shifted the criterion toward the liberal side, in line with previous findings^22, 24, 25, 42^. Dovetailing this attentional enhancement of conscious report effect in behavior, we identified three neural clusters in which validly cued seen targets elicited stronger neural activity than invalidly cued ones. This enhancement occurred first in high-level visual areas as a fast transient target-related activity, then in FP and temporal regions showing a sustained activity, and finally in bilateral PFC, presenting late accumulation activity, which lasted until the motor response.

Previous evidence showed attention modulates neural responses across the visual cortical hierarchy with an increasing magnitude from early to higher-level visual areas^43, 44^, but the precise location of its interaction with consciousness in the visual cortex was little known. Our results clarified that spatial cuing first potentiated awareness-related activity in the high-level visual areas (Visual cluster; fusiform gyrus, middle temporal gyrus, inferior temporal gyrus and inferior parietal cortex), but not in the early visual areas.

The Sustained cluster included hotspots around the right superior frontal gyrus and inferior parietal lobule which were connected by the SLF II network. The attentional enhancement of conscious perception in this cluster may reflect recurrent neural activity which provides a specific neural substrate for neural amplification as suggested by the GNWH^10^. The involvement of the SLF II in the attentional modulation of conscious perception is in line with previous clinical evidence showing that damage to the right SLF II is the best anatomical predictor of the occurrence of neglect signs in stroke patients^28, 45^. Importantly, transitory electrical inactivation of the SLF II in the human right hemisphere in a patient undergoing brain surgery provoked severe, if transient, rightward shifts in line bisection, akin to signs of left spatial neglect^46^. The present evidence specifies the temporal dynamics of the right hemisphere SLF II network, by demonstrating its role in attentional amplification of future targets.

The Late accumulation cluster included focal contacts around the right mesio-frontal and supplementary motor areas connected by SLF I-III network. This cluster showed a late accumulation activity until report, which was enhanced by spatial cueing. This interaction is consistent with its known role of SLF I network in evidence integration and decision making^47^. Alternatively, the activity of this cluster might reflect temporal expectancy for conscious report of targets^48, 49^. In line with early EEG studies^50^, our results demonstrate that attention could enhance conscious expectancy through this FP network.

A reversed pattern of interaction between attention and conscious report occurred when targets appeared at the uncued location: when reported, these targets elicited early, sustained activity in the right hemisphere TPJ and IFG (reorienting network)^51^, connected by the SLF III. The activity of this Reorienting cluster supports the role of the SLF III network in the conscious perception of targets preceded by an invalid cue at the “wrong” location. Previous neuroimaging evidence has shown the implication of this network in reorienting attention from the invalidly cued location to the targets occurring on the opposite side^26^. Importantly, our time-resolved results specify that this activity happens earlier than previously thought, before target presentation. This early activity (pre-target until around 150ms post-target) suggests an anticipatory (“lookout”) activity for unexpected events such as invalidly cued targets and if necessary, it reorients attention before the attentional enhancement implemented by the SLF II network. Consistent with this hypothesis, early SLF III network activity decreased when the reported targets appeared at the validly cued location.

Finally, independent of attention, seen, but not unseen, targets elicited late sustained activity in the Conscious report cluster located around the posterior portion of left dorsolateral PFC. In line with previous neuroimaging evidence^52^, this neural activity might reflect the integration of sensory evidence and the formation of decision variables in post-orienting processes that are closely associated with conscious access mechanisms. Another possibility is that the left dorsolateral PFC biases perceptual decisions in conditions of uncertainty^53^ such as near-threshold detection. It remains to be seen how our intracerebral data relate to putative markers of consciousness derived from surface EEG^16^.

Our RNN model of the task displayed striking similarities with the human intracerebral data, and allowed us to make causal inferences on the inter-network interactions of the neural clusters we observed. Two distinct components of the attention/consciousness interactions, attentional enhancement and attentional reorienting, clearly emerged in the trained RNN models. The Sustained excitatory units receive the majority of stimuli input and selectively enhance target-related information for conscious report. The Reorienting excitatory units directly receive a smaller branch of visual input. The inhibitory units in the Sustained and Late accumulation RNN clusters receive excitatory input from the Reorienting units. Critically, these results suggest that the right Reorienting neural cluster reorients spatial attention by monitoring the unexpected events in the environment and stopping the ongoing stimulus processing by inhibiting inappropriate activities of other clusters. RNN lesion data further support specific causal contributions of the identified neural clusters and their associated SLF networks in modulating conscious report of near-threshold stimuli. Our model predictions are consistent with neuroimaging evidence showing the involvement of right-hemisphere FP networks in attentional enhancement^54^ and inhibitory control^7, 55^. Additionally, a previous simulation study of physiological leftward bias (pseudoneglect) found a similar excitatory influence of the right ventral attentional network linked by the SLF III on the dorsal attentional network^56^. Our results extended this prediction to an excitatory-inhibitory neuronal interaction with conscious report, at least for detecting near-threshold stimuli. Strong inter-regional excitation, balanced by local inhibition, can enable reliable sensory signal propagation to the PFC, which in turn can lead to “global ignition”^57^ and pave the path to conscious visual processing, consistent with the GNWH^58^. More direct causal evidence of the interplay of these FP attentional networks for conscious perception comes from the finding that damage to the right dorsal PFC and decreased microstructural integrity of the SLF III impaired the conscious perception of near-threshold information^59^.

Taken together, the observed interactions between exogenous attention and conscious report provide new, compelling evidence that support current models of consciousness, which explicitly conceptualize the role of attention in consciousness, such as the gateway hypothesis^9^, and the GNWH^11^. Our findings specify the temporal dynamics and computational mechanisms underlying attention and consciousness interactions. More generally, our results shed light on the role of dorsolateral, ventrolateral and orbital PFC as well as high-level visual cortex, in human conscious report of near-threshold targets. Previous work on non-human primates showed that reported stimuli were associated with strong sustained PFC activity^60^. However, this late extensive PFC activity might reflect decision-making instead of conscious experience^15, 16^. Our findings reconcile this debate by demonstrating, on the one hand, the late accumulation activity in right FP areas connected by SLF I-III, which lasted until target report; on the other hand, our results also show the sustained activity for consciously reported target in left PFC and right FP areas connected by SLF II-III, independent of decision time.

The present study has some limitations. First, the use of a subjective detection task to probe conscious perception, in which participants had to choose between 3 alternatives indicating the perceived location of the target (left or right) or its absence. This measure provides information concerning whether participants consciously perceived the target, but it does not provide more subtle information regarding the participants’ awareness of their perception, unlike confidence ratings^61^. Future research might also use no-report paradigms^62^ to dissociate awareness-related neural activation from potential decision-making effects. Second, patients were instructed to maintain central fixation, but the clinical setting did not allow us to use eye-tracking recording. Third, cortical coverage was necessarily limited (e.g in the superior parietal cortex), because it was obviously only determined by clinical needs. Fourth, our subjects had chronic epilepsy, although contacts with epileptic activity were excluded from analysis. Last, we didn’t report data in subcortical routes whose neural activities parallel cortico-cortical networks. Recent work has shown that the thalamus is a major hub for both consciousness and attention, and the recurrent thalamocortical loops might be a major feature in supporting consciousness^63–66^.

Despite these considerations, the present evidence establishes the neural dynamics of distinct FP networks and high-level visual areas in the attentional modulation of conscious report of near-threshold stimuli, which is one of the hallmark concepts distinguishing different consciousness theories^5, 8^. Our findings establish specific roles for the right-hemisphere SLF II network in the attentional enhancement of near-threshold targets, for the right-hemisphere SLF III network in perceiving previously unattended targets, and confirms the hypothesized role of left-hemisphere dorsolateral PFC in perceptual decision. This attention/consciousness interaction relies on specific excitatory and inhibitory inter-network interaction mechanisms that causally contribute to conscious perception of near-threshold targets. Thus, distinct, hemisphere-asymmetric fronto-parietal networks support attentional gain and reorienting in shaping human conscious experience.

## Acknowledgements

This work is supported by the Agence Nationale de la Recherche through ANR-16-CE37-0005 527 and ANR-10-IAIHU-06, by the Fondation pour la Recherche sur les AVC through FR-AVC-017, 528 by Fondation Assitance Publique-Ho_pitaux de Paris (EPIRES-Marie Laure PLV 529 Merchandising), and by funding from Dassault Systèmes.

## Author contributions

Conceptualization: DJB, PB

Data curation: KL

Formal analysis: JL, DJB

Methodology: JL, JDS, DJB, PB, TSM

Visualization: JL

Funding acquisition: PB, TSM, VN, CA, JL

Project administration: PB

Resources: KL, SFV, VN, CA, VL

Software: JL, DJB, JDS, SFV, TSM

Supervision: TSM, PB

Writing - original draft: JL

Writing - review & editing: All authors

## Competing interests

Authors declare that they have no competing interests.

## Materials & Correspondence

Requests for materials and correspondence should be addressed to Jianghao Liu, Tal Seidel Malkinson and Paolo Bartolomeo.

## Methods

### Participants and intracerebral recordings

We recruited sixteen patients who underwent presurgical evaluation of pharmaco-resistant focal epilepsy with intracerebral encephalography (iEEG) implantation, at the Department of Neurosurgery of the Hôpital Pitié-Salpêtrière, Paris, France. All participants had normal or corrected-to-normal vision (age mean ± SD: 35.0 ± 8.2; 10 women; 14 right-handed) and provided their written informed consent (CPP Paris VI, INSERM C11-16 (2012-2020); CPP INSERM C19-55). Three patients were excluded from the data analysis due to poor data quality (two patients had corrupted neural data files and one patient had response times inferior to 150 ms in 32% of trials), leaving a total of thirteen patients in the final sample (age mean ± SD: 34.7 ± 8.7; 7 women; 11 right-handed). Patients were implanted with 4-13 multilead stereotactic depth electrodes (AdTech®, Wisconsin) endowing 4-12 platinum contacts with a diameter of 1.12 mm and length of 2.41 mm, with nickel-chromium wiring. The distance between the centers of two contiguous contacts was 5 mm. The location of electrode implantation was based exclusively on clinical criteria. In five patients the neural activity was recorded with a 128 channels clinical video-EEG recording system (SD LTM 64 BS, Micromed® S.p.A., Italy), sampling at 1024 Hz with a band-pass filter of 0.15 Hz to 250 Hz. In the other eight patients, the recording was implemented with a Neuralynx system (ATLAS, Neuralynx®, Inc., Bozeman, MO) which allowed recording up to 160 depth-EEG channels sampled at 4 kHz with 0.1Hz to 1000 Hz band-pass. The patient-dependent least active contact, preferably in the white matter, was selected as the reference electrode.

### Experimental task

The stimuli presentation was controlled by E-Prime 2.0 software (SCR_009567) running on a laptop with a 60 Hz refresh rate. Three black boxes (4.9° long and 3.6° large) were arranged around a central fixation point, persisted for the whole duration of the trial, with 6° horizontally separating the central box center from the peripheral boxes’ center. The two peripheral boxes were located in the lower visual field, 4° of visual angle under the central box. Fig.1A illustrates the experimental procedure. Participants were instructed to fixate their gaze at the central fixation cross throughout the test and to respond as fast and accurately as possible. Following the appearance of the fixation and the three placeholder boxes for 1,000-1,500ms, a peripheral cue consisting of a black dot (1° diameter) was presented for 50ms at the upper external corner of one of the two peripheral boxes. Three-hundred ms after the visual cue onset, a target stimulus was presented for 16ms in one of the two peripheral boxes (or not presented in “catch” trials). The target stimuli were tilted Gabor patches with a spatial frequency of 5 cycles and the diameter of 2.5° visual angle, chosen among 12 equally spaced between 0 and 180°, excluding vertical and horizontal orientations. After a 484ms delay from the target offset, participants performed a 2-alternative forced choice discrimination, indicating the direction of the tilt among two possibilities presented on the screen and distant by 30° from one another (discrimination task). Then a response screen appeared prompting the participants to report the perceived presence or absence of the Gabors (detection task), by selecting one of two opposing arrows (indicating the perceived location of the target), or the word ‘absent’ under the arrows. Trials lasted until participant’s response or for a maximum of three seconds. Participants performed eight recording blocks, each consisted of 110 randomized trials including 88 target-present trials and 22 target-absent “catch” trials. Participants were informed that cues were non-predictive, i.e. in target-present trials cues indicated the target location in 50% of trials (validly cued) and the opposite location on the remaining 50% of the trials (invalidly cued). Before the recording blocks, participants performed a target contrast calibration session in order to estimate the individual perceptual threshold contrast for 50% seen targets. The calibration consisted of two randomly interleaved, one-up one-down staircases, converging toward a detection rate of 50%. Staircase stimuli were the same as the main paradigm across all conditions.

### Behavioral analysis

For each participant, we first excluded trials with response time (RT) faster than 150 ms or exceeding three standard deviations. Statistical tests in behavioral analysis were performed using JASP software (version 0.16.0.0, https://jasp-stats.org/). We performed a two-way ANOVA with the factors of cue validity and conscious report (in the detection task) on the discrimination accuracy. RTs analysis was not reported since participants had to wait for the response screen to give their responses.

Using a nonparametric measure^67^, we conducted Signal Detection Theory (SDT) analysis^31^ to evaluate the bias produced by the cue validity on participants’ perceptual sensitivity *a’*. We computed the mean percentage of seen targets when the Gabor was presented (Hits) and when the Gabor was absent (false alarms; FA). The criterion *C* summarizes the distance of the threshold relative to the noise distribution from the threshold of an ideal observer. A smaller value of *C* represents a more liberal threshold in target detection.

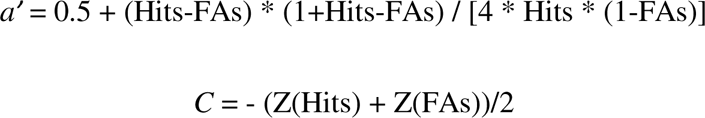

We compared detection rate, false alarm rate, sensitivity, and criterion between valid and invalid trials with paired sample *t-*test.

### iEEG preprocessing

Spatial localization of each electrode was recovered using the Epiloc toolbox^68^ developed by the STIM engineering platform in the Paris Brain Institute (https://icm-institute.org/en/cenir-stim) with co-registered pre-implantation 1.5T or 3T MRI scans and post-implantation CT and MRI scans. After the normalization of MRI-pre, MRI-post and CT-post into the MNI space, contact localization was automatically labeled referring to Desikan-Killiany-Tourville atlas parcellation^69^ in the patient’s native space, using Freesurfer image analysis suite (https://surfer.nmr.mgh.harvard.edu/) embedded in the Epiloc toolbox, followed by a manual verification and correction, if necessary. Fig. 1C and Table. S1 show the localization of usable contacts referring to Desikan-Killiany-Tourville atlas parcellation ^69^: 29 contacts (4%) from occipital, 332 (46%) from temporal, 78 (11%) from parietal, 202 (28%) from frontal, 24 (3%) subcortical and 62 (9%) in white matter. There were 288 contacts in the left hemisphere and 439 contacts in the right hemisphere.

Signal preprocessing was conducted using Matlab (R2018b, The MathWorks, Inc.) and FieldTrip toolbox^70^. First, all signals were down sampled to 512 Hz and all contacts were re-referenced to their adjacent neighbor contact on the same electrode, yielding a bipolar montage, in order to ensure that iEEG signals could be considered as originating from a cortical volume centered within the two contacts. Coordinates of bipolar contacts were computed as the mean of the MNI coordinates of two adjacent contacts composing the bipole. An initial visual inspection of continuous signals was performed to remove time segments showing transient epileptic or interictal activity. Contacts with excessive epileptic spikes or near suspected epileptic focus were also rejected. We extracted time courses from −1,300 to 1,200ms around target onset for trial epochs. A second artifact inspection was performed on the epoched data, where trials and contacts with excessive maximal signal, z-value, variance or kurtosis of the signal distribution were rejected. After signal preprocessing, 727 of the bipolar contacts out of 887 contacts were retained for further analysis.

We then adopted a pseudo-whole-brain analysis approach, by pooling contacts across all thirteen patients on a standardized brain in MNI space. High-frequency broadband (HFBB) power (70-140Hz), a proxy of spiking activity of the local neuronal ensemble^32, 33^, but also see^34^, was extracted from each bipolar contact by wavelet time frequency transformation using the Morlet wavelets implemented in Fieldtrip (ft_freqanalysis), in fourteen equally spaced center frequency bands. We kept high-frequency band power time courses from −800 to 1,000ms to target onset to discard the 1/f signal drop off at the edges. Baseline normalization was applied on each trial by means of a z-score relative to the period in the 200ms prior to cue onset. Finally, HFBB powers were down-sampled to 100Hz for further analysis.

### Temporal embedding visualization with t-SNE

We visualized neural activities and unit activities in a two-component space by a machine learning visualization approach, t-distributed stochastic neighbor embedding (t-SNE)^71^. In neural data, we averaged neural activities across contacts by conditions and time points, of which the resulting matrix served as input to compute temporal embedding. t-SNE was computed by an implementation in Scikit learn 1.0.2 in Python 3. We adopted a perplexity of 30 and a learning rate of 100. The embedding was initiated with the PCA option and optimized upon 1000 iterations, by default. The neural data in each condition was projected to a two-dimensional t-SNE embedding. Thus, the ensemble of time points formed a temporal trajectory of neural activities in a reduced manifold. A larger difference between two trajectory components represented a more distinct neural activity pattern between the two conditions. In RNN simulation, we averaged unit activity across units by condition and time point, and kept the above parameters in computing its t-SNE temporal embedding.

### Trajectory k-means clustering

We applied a novel clustering approach based on *k*-means clustering to classify contacts by their temporal profiles^35^, implemented through Matlab (R2018b, The MathWorks, Inc.). This data-driven approach was able to capture the prototypical patterns of neural dynamics that might be sensitive to cue validity and seen/unseen reports. We conducted clustering on all bipolar contacts. In each contact, we took the trajectories of the mean target-locked activity across an 8-dimensional condition space (target side, cue validity and seen/unseen report). Activity across conditions was z-scored relative to the distribution of the trials’ entire duration. Contacts were then iteratively partitioned (10000 iterations) into 2-12 clusters, in which each contact was assigned to the cluster with the nearest centroid trajectory. This was achieved by minimizing the sum of time-point-by-time-point Manhattan distances across conditions, to quantify trajectories similarity while preserving temporal order. Based on the silhouette evaluation (*silhouette* in Matlab), we adopted a ten clusters solution, which reflected the highest average silhouette score (Fig.S2A). The partition of the 13 patients’ contacts to clusters is shown in Fig.S2B. In order to identify the consistency of clusters across different numbers of clusters K, we inspected clusters’ trajectory profiles in each number of clusters. We plotted the trajectories of five clusters that showed significant consciousness effects in the K =10 cluster solution (Fig.S2D). The minimal variation of the number of contacts in each cluster demonstrated the stability of the contacts in the five selected clusters across k-means solutions (Fig.S2C).

### Conscious report and interaction-related neural activity

To explore how our experimental manipulation of attention and consciousness influenced the clusters’ neural activity, we performed time-resolved three-way ANOVAs with the factors of target side (left/right), cue validity (valid/invalid), and conscious report (seen/unseen). We tested on HFBB power in both cue-target period (from −300ms to target) and post-target period (from target onset to 500ms), across contacts on each cluster. For each contact, we averaged HFBB power across trials by conditions. Holm-Bonferroni correction was applied over all the time points for multiple comparisons. For clusters showing significant consciousness effect, post-hoc comparisons were performed on time points where the Cue validity x Conscious report interaction was significant, with Holm-Bonferroni correction. Further, in order to compare the degree of attentional enhancement across clusters that showed higher neural activity for validly cued seen targets, we performed a linear contrast testing around time points where the clusters showed significant effects.

### Response time (RT) modulation of neural activity

To understand the functional roles of neural clusters, we related the neural activity to the response time. In each cluster, we pooled trials from all conditions and contacts. We then sorted the trials and binned them into 20 quantiles according to their response time in the discrimination task. We compared RT-bins neural activity using time-resolved one-way ANOVA in a short time window (target onset to 1,000ms, to avoid the influence by the neural activity of the subsequent trial). To specifically test whether higher sustained neural activity was related to the slower response time, we then compared the neural activity of the 10 RT bins with slowest response to the neural activity of the 10 RT bins with fastest response. Holm-Bonferroni correction was applied over all the time points. We also visually compared the RT bins sorted neural activity in a longer time window (target onset to 5,000ms) in order to identify the neural activity patterns associated with visual modulation and task report in each cluster.

### White matter tracts computing

We dissected white matter tracts connecting parietal contacts with frontal contacts. We modeled each intracerebral contact by a 3 mm-sphere. In each cluster, we created two region-of-interests (ROIs), respectively consisting of frontal contacts and parietal contacts. The parcellation of contacts was referred to Desikan-Killiany-Tourville atlas (see details in iEEG preprocessing and Table S1 for contact numbers). We used a dataset that includes 176 preprocessed healthy individuals tractography acquired at 7 tesla by the Human Connectome Project team ^37^. We performed tracts filtering in TrackVis toolbox ^72^ to obtain tracts connecting frontal contacts and parietal contacts in frontal association tracts ^38^ (three branches of the superior longitudinal fasciculus; uncinate; long and anterior segments of the arcuate fasciculus; inferior fronto-occipital fasciculus). For each cluster, the tractography maps of the 176 healthy individuals were subsequently binarised and then smoothed with a three-dimensional Gaussian filter (full width at half-maximum was 5 mm, equivalent to a sigma of 2.123). To test the presence of tracts across individuals, we used a threshold-free cluster enhancement (TFCE)-based non-parametric test, with 1000 permutations (*“randomize”* function in FSL) and a height threshold of 0.95 to control significance level at *p*<0.05. We computed the volumetric ratio of the labeled tracts in each cluster with those of the standard atlas in BCBtoolkit ^73^ (http://toolkit.bcblab.com/), where we filtered the atlas probabilistic maps with a threshold of 80% to reduce their overlapping.

### Task-optimized recurrent neural network model

Recurrent neural networks (RNNs) are networks in which neurons (units) can send and receive feedback to and from each other. Therefore, the activity of the units is affected not only by the current external stimulus, but also by the current state of the network^74^, which makes RNNs ideally suited for computations that unfold over time such as holding the information of cue position or accumulating target-related evidence for decision making. If trained RNN accomplishes the behavioral task with a performance comparable to the human ones, the RNN hidden unit activities could provide unique insight about its computations in task representation. Moreover, RNN unit activities might also appropriately predict neural processing^75, 76^.

Our RNN model contained a single layer trained with mini-batch gradient descent learned by backpropagation. Before time discretization, the network activity follows a continuous dynamical equation:

where, and denote the input, recurrent state, and output vectors, respectively., are the connection weight matrices of the input layer (a *2* x *N* matrix), the recurrent layer (*N* x *N*) and the output layer (*N* x *2*). and are constant biases into the recurrent and output units. The network is time-discretized with positive activity. is the simulation time-step and is an intrinsic timescale of recurrent units which was set to 100ms. denotes the independent Gaussian white noise processes with zero mean and unit variance, and is the strength of the noise. is a nonlinear transfer function, which was set as a rectified linear activation function (ReLU).

Similar to previous studies training RNNs to perform cognitive tasks^77–79^, we abstracted the relevant visual stimuli properties from the patient task (Fig.1A), rather than feeding the exact same visual inputs to the RNNs. Specifically, visual stimuli from the left and the right fields of view were modeled as two separate noisy inputs (Fig.4B). The magnitude of the background visual noise along the trial was set as Gaussian noise of the mean of zero and standard deviation (SD) of 0.05. The task began with a fixation period of 200ms followed by a visual cue randomly presented at either the left or the right side. A target, separated from the cue by a fixed delay period of 300ms, was then presented on the cued position (validly cued) or on the opposite side (invalidly cued) with equal probability, or absent in 20% of “catch” trials. After a second fixed delay, the network produced two outputs, each ranging from zero to one, to respond whether there was a target detected. Similar to the human task, RNN outputs were combined to calculate a discrimination and a detection performance. The discrimination accuracy is the ratio of correct response with forced-choice in distinguishing target side when the target were presented. The detection rate equals the ratio of trials in which the network made the correct response about the target presence or absence.

We implemented the model training with PsychRNN^80^, a toolbox backended by TensorFlow. We adopted the default setting of the package regarding the regularizers (i.e. penalties added to prevent over-fitting to the training data), weight initializer (i.e. by a glorot normal distribution from −1 to 1) and the loss function (i.e. mean squared error). The connections between hidden units were constrained according to Dale’s principle: neurotransmitters tend to be either excitatory or inhibitory such that the post-synaptic weights of each recurrent unit are all of the same sign^79^. There were 80% of units fixed to be excitatory and the remaining 20% of units were inhibitory. The strong inhibitory signaling in the recurrent neural network enables stable temporal dynamics^78^. The cue contrast was set as 0.30 standard deviation (SD) and we trained RNNs, one at a time, with various target stimulus levels from zero to 0.13 SD, which mimics the near-threshold targets setting in the patient task. The RNNs were trained with 150,000 iterations and the model accuracy was tested on 50 batches of 50 sample trials for each target contrast, respectively. Finally, we fitted a psychometric detection curve by logistic function in the detection task and in the discrimination task, respectively. To verify the validity of the model, we compared the performance of trained models with an untrained model that was initialized with random Gaussian connectivity weights without feeding stimuli inputs for learning. We also tested RNNs with different hidden units’ size (128, 50, 32 and 16) and found out that those RNNs could not achieve human-level performance when the units were less than 16. To keep sufficient units for further clustering analysis, we adopted the number of hidden units N as 50 to balance the model’s complexity and variability.

### Trajectory k-means clustering on RNN unit activities

We applied the above-mentioned trajectory clustering method to classify dynamic patterns of the fifty hidden units for intermediate target contrast (0.10 SD). We generated 20 batches of 50 trials (1000 trials in total) and averaged unit activity by condition (all conditions: validly cued seen, invalidly cued seen and no target). Unit trajectories were iteratively partitioned into 2-10 clusters across the three-dimensional condition space. The highest averaged silhouette value was obtained while the number of clusters equals five (Fig. S4D). We compared seen/unseen trials by time-resolved *t*-test in the post-target period (target onset to 1,000ms, see Fig. 4D) which could include information after output responses. Holm-Bonferroni correction was applied over all the time points. We also conducted the same *k*-means clustering analysis on a reduced target contrast (0.03 and 0.06 SD). However, only a late cluster showed a significant consciousness effect. It might be due to the low target contrast that other clusters did not attend to significance level.

### Computing similarity between neural clusters and RNN clusters

In the four neural clusters showing the cue modulation on conscious perception (Visual, Sustained, Late accumulation, and Reorienting), we averaged neural activity in seen and in unseen trials from cue onsets to 500ms post-targets. We then generated 500 batches of 50 trials with the RNN model simulation for both trained and untrained models. In each batch, we averaged RNN clusters’ unit activity in seen and in unseen trials. The similarity was quantified by Pearson correlation coefficient between the RNN clusters’ temporal trajectory to the neural cluster ones, with the averaged coefficient of conditions. Therefore, we obtained a distribution of correlation coefficients with 500 samples for the trained and for the untrained models. Finally, we conducted a one-sided permutation test with 1,000 permutations to compare the two distributions, in order to identify the RNN clusters being significantly more similar to the neural clusters in the trained model than in the untrained model.

### Directed connection weight graph

The iEEG contacts of the neural clusters were pooled from different patients, which limits the possibility to analyze their inter-cluster connections. However, the modeling provides the possibility to examine how unit clusters are inter-connected as an integrated model. To this end, we extracted unit input weights, directed unit-to-unit connection weights and output weights of the trained model (Fig. S4E). In each cluster, we observed both excitatory (E) and inhibitory (I) type units, of which the number was fully task-optimized without any prior. The Sustained cluster contained 16 E and 4 I units. The Late accumulation cluster had 6 E and 2 I units. The Reorienting cluster had 8 E and 3 I units. We grouped units of the same E/I type in each cluster, resulting in six groups in total. We then computed a directed cluster connection graph by averaging unit connection weights from one group (“pre-synaptic”) to each of the other groups (“post-synaptic”). For example, to compute the connection weight from the Sustained excitatory group (16 units) to the Reorienting excitatory group (8 units), we averaged a total 128 directed unit-to-unit connection weights. One-sample *t*-test was performed with significance level controlled at *p* < 0.05, Holm-Sidak corrected for multi-comparison among groups of units.

### Lesion analysis

To confirm the functional roles of the clusters, we lesioned units, one cluster at a time, and monitored the decrease in task performance. This was achieved by setting unit’s connection weights with inputs, all recurrent units and output to zero. We tested the task performance of the lesioned models with generated 50 batches of 50 trials.

## Supplemental information

**Fig S1.**
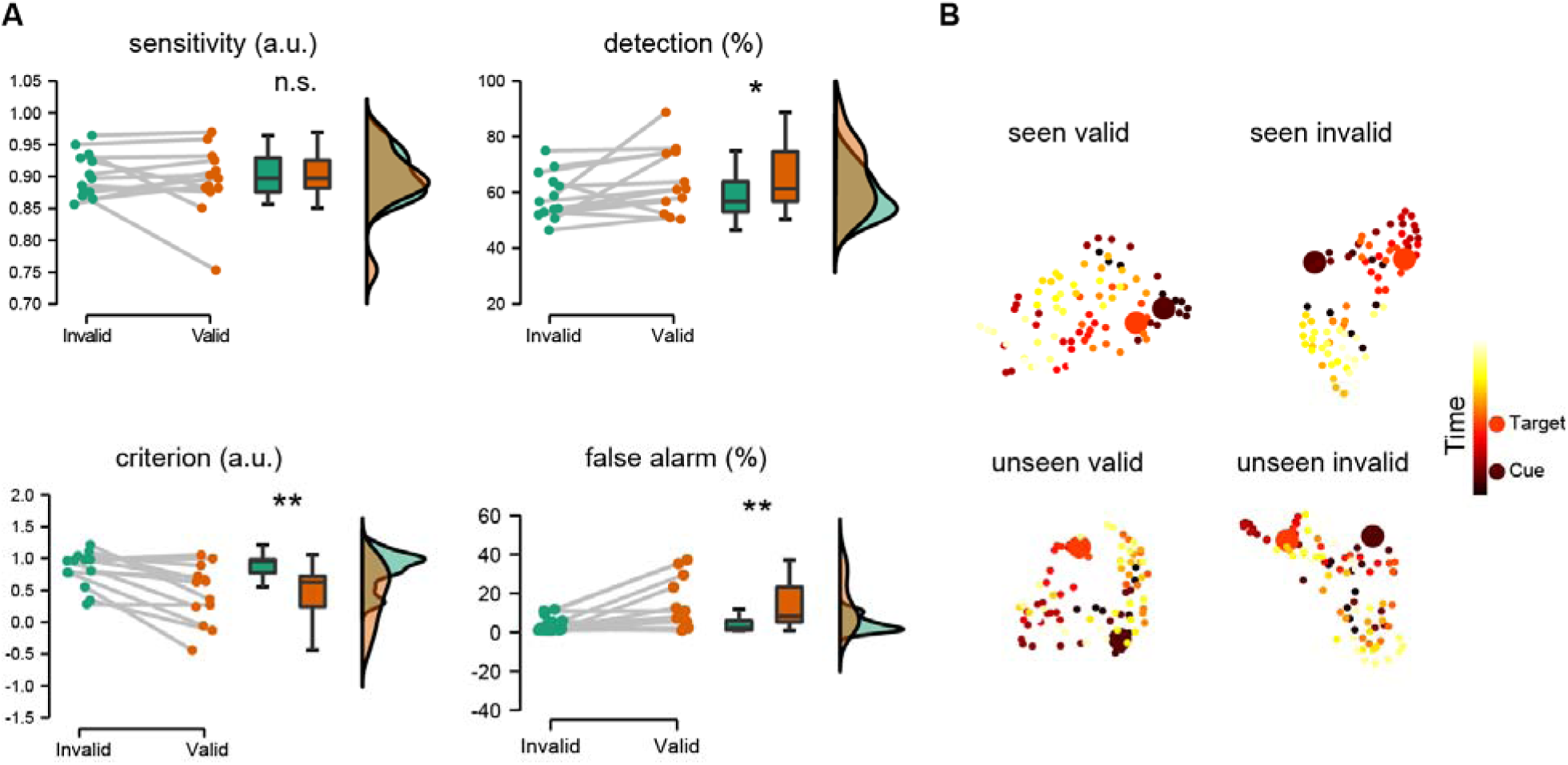
Signal detection theory analysis and visualization of neural activity components. **A.** Comparison of sensitivity, detection rate, criterion, and false alarm rate in valid *versus* invalid trials. Dots represent individual performance. \**p*<0.05; \*\**p*<0.01; n.s.: not significant; a.u. arbitrary unit. **B.** Two-dimensional t-distributed stochastic neighbor embedding (t-SNE) visualization of neural activity components of all contacts. Color map represents time points, darker for earlier and lighter for later time points.

**Fig S2.**
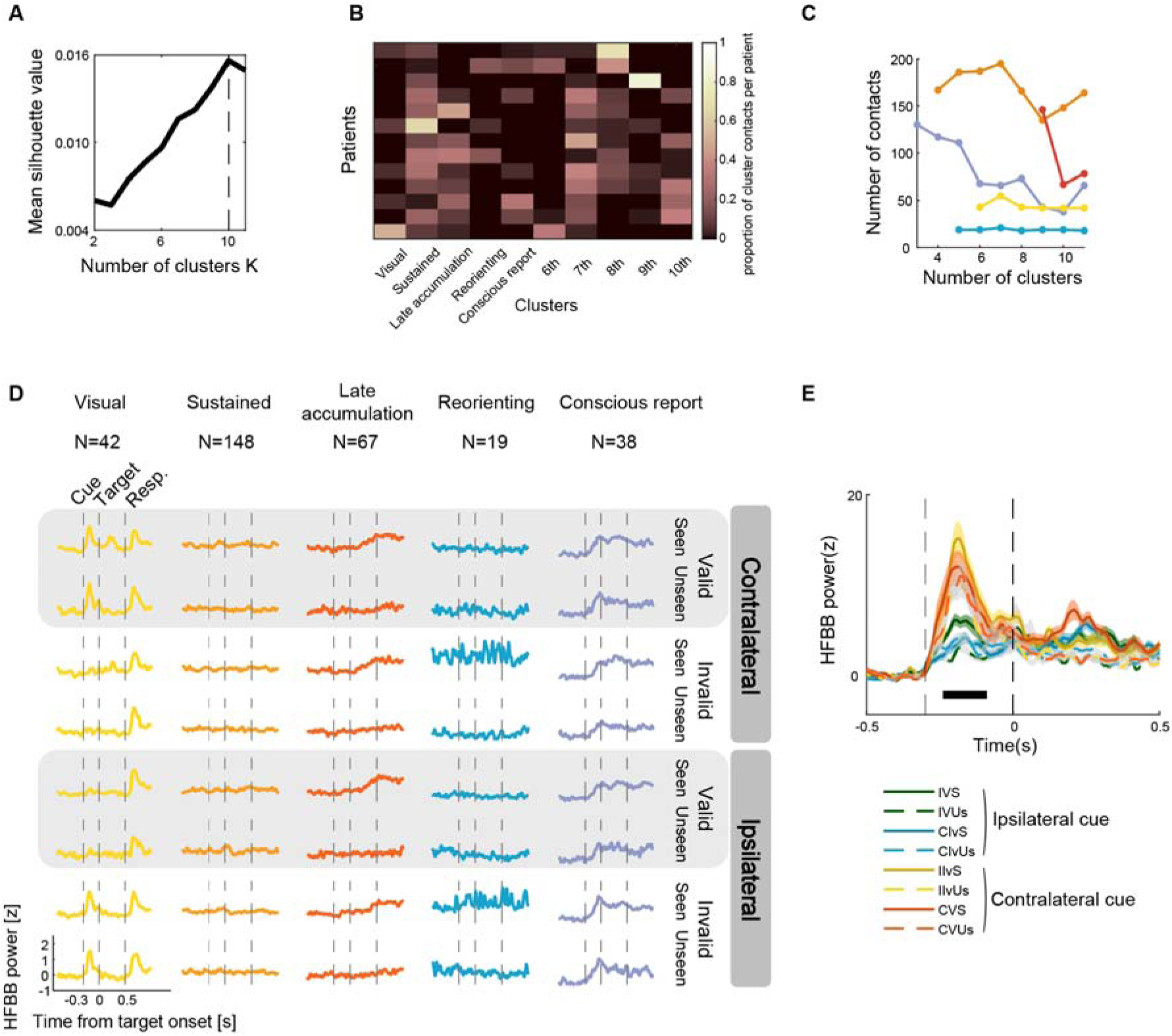
Clustering the neural activities of intracerebral contacts. **A.** Mean silhouette value of all contacts across k-means solutions. **B.** Partition of contacts by cluster for patients. Rows represent patients and color map denotes the proportion of contacts in each of the clusters per patient. This demonstrates that clusters did not result from any single participant’s trajectory activity, but rather reflected temporal patterns across many participants. **C.** Cluster stability. The change of the number of contacts across *k*-means solutions. **D.** Visualization of trajectory neural activities by conditions for k-means solution K = 10. Resp. -response screen, after which the participants were allowed to respond. N -number of contacts in the cluster. **E.** Neural activity in the Visual cluster. During the cueing period (−300ms to 0), the Visual cluster showed a target side x validity interaction, with higher neural activity for contralateral cues than ipsilateral cues. Black horizontal bar for all *p* < 0.05, Holm-Bonferroni corrected. I/C: Ipsilateral or Contralateral target; V/Iv: Valid or Invalid; S/Un: Seen or Unseen.

**Fig S3.**
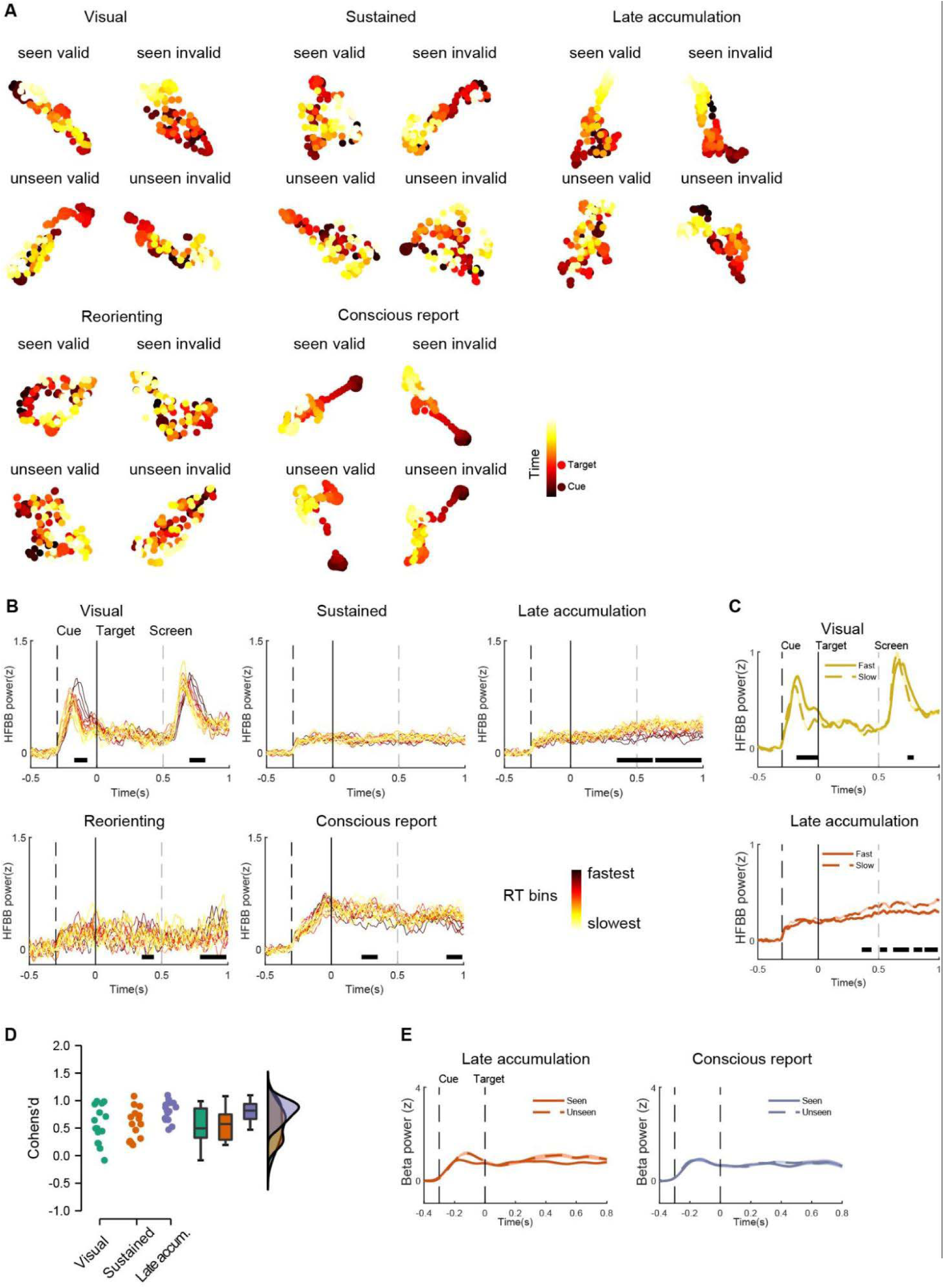
Intracerebral clusters neural activity. **A.** Neural component visualization with t-SNE. Colormap represents time and dots represent neural activity per time point, darker for earlier and lighter for later time points. **B.** Visual stimuli- and RT-modulation of target-locked neural activity in a time window from target onset to 1,000ms. In each cluster, the trials in all conditions were divided into 20 quantile RT bins of equal probability. RT bins were sorted according to their mean RT in the discrimination task from fastest (in yellow) to slowest (in red), with neural activity pooled across contacts in each cluster, and RT-related modulation was tested using a time-resolved one-way ANOVA, performed on the neural activity across RT bins. Black horizontal bars indicate *p* < 0.05, Holm-Bonferroni corrected. **C.** Comparison of neural activities of the ten fastest and the ten slowest RT-bins by time-resolved *t*-test. Black horizontal bar for all *p* < 0.05, with Holm-Bonferroni correction. No significant difference was found in the Sustained, Reorienting and Conscious report clusters. **D**. Linear contrast comparison of the attentional enhancement effect across the Visual, Sustained, and Late accumulation clusters. Cohen’s d was derived from time-resolved *t*-test for the contrast (seen valid - unseen valid) - (seen invalid - unseen invalid) around the significant interaction time point in each cluster. Late accum. = Late accumulation cluster. Dots represent Cohen’s d values at different time points. **E.** Beta band power. No decreased beta activity was observed in the Late accumulation and the Conscious report clusters, indicating the activity in these clusters does not reflect motor planning.

**Fig S4.**
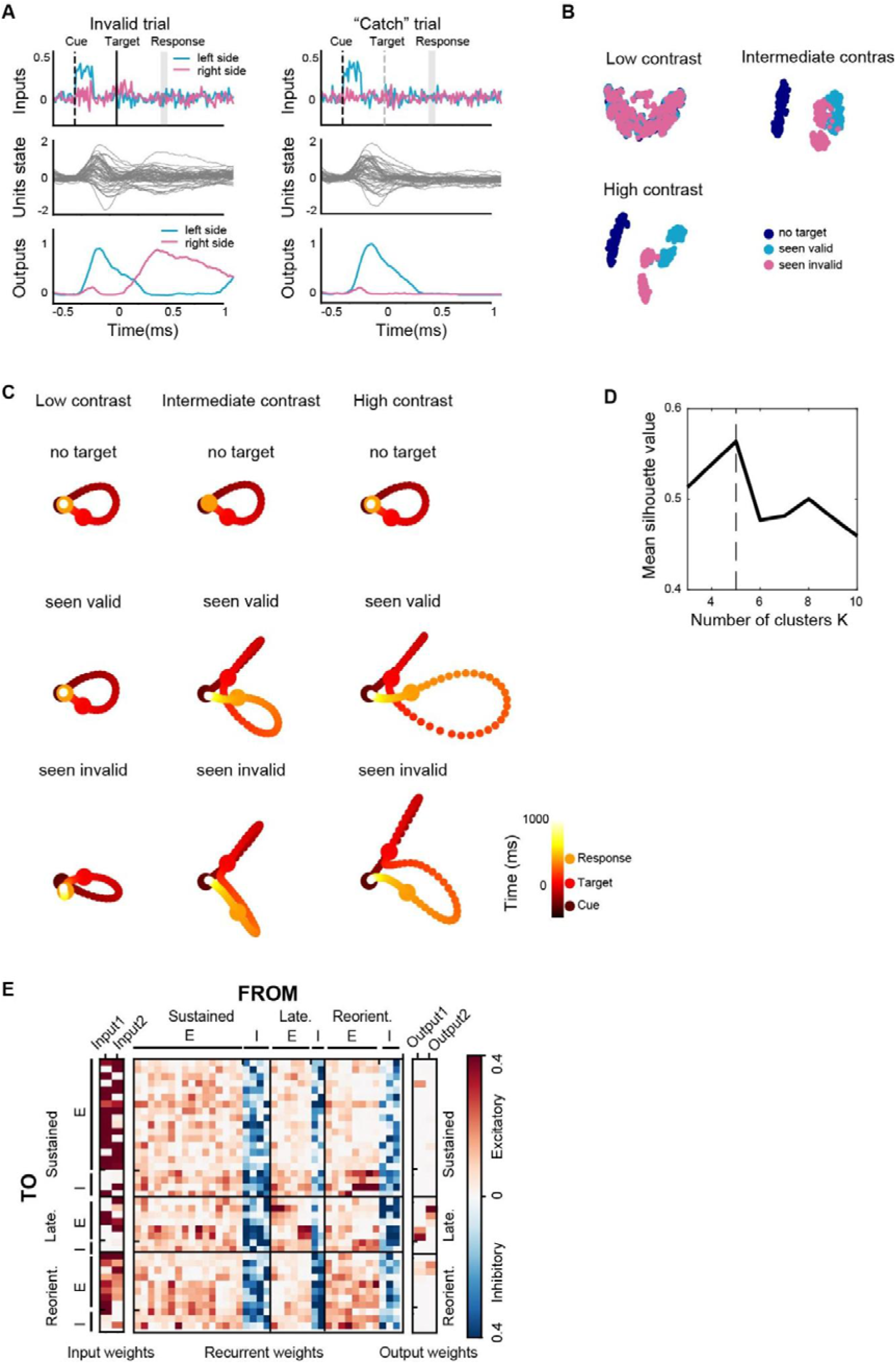
Task representation and unit activity in the recurrent neural network model. **A.** Example of task structure in invalid (left panel) and in target-absent “catch” trials (right panel). **B.** Two-dimensional t-SNE visualization of unit activity of all units, for low, intermediate and high target contrast levels, respectively. Dots represent trials. **C.** Two-dimensional t-SNE visualization of unit components of all units. Colormap represents time points. **D.** Mean silhouette values of all units across k-means solutions in trajectory-clustering of RNN units. **E.** Directed connection weights of the task-optimized trained model. Left columns: input weights mediating sensory enhancement gain; Middle columns: directed unit-to-unit connection weights; Right columns: output weights mediating report gain). Connections go from unit columns (“pre-synaptic”) to unit rows (“post-synaptic”). Each cluster contains both excitatory (E) and inhibitory (I) units. Red = excitatory weights; blue = inhibitory weights; Late. = Late accumulation; Reorient. = Reorienting.

**Fig S5.**
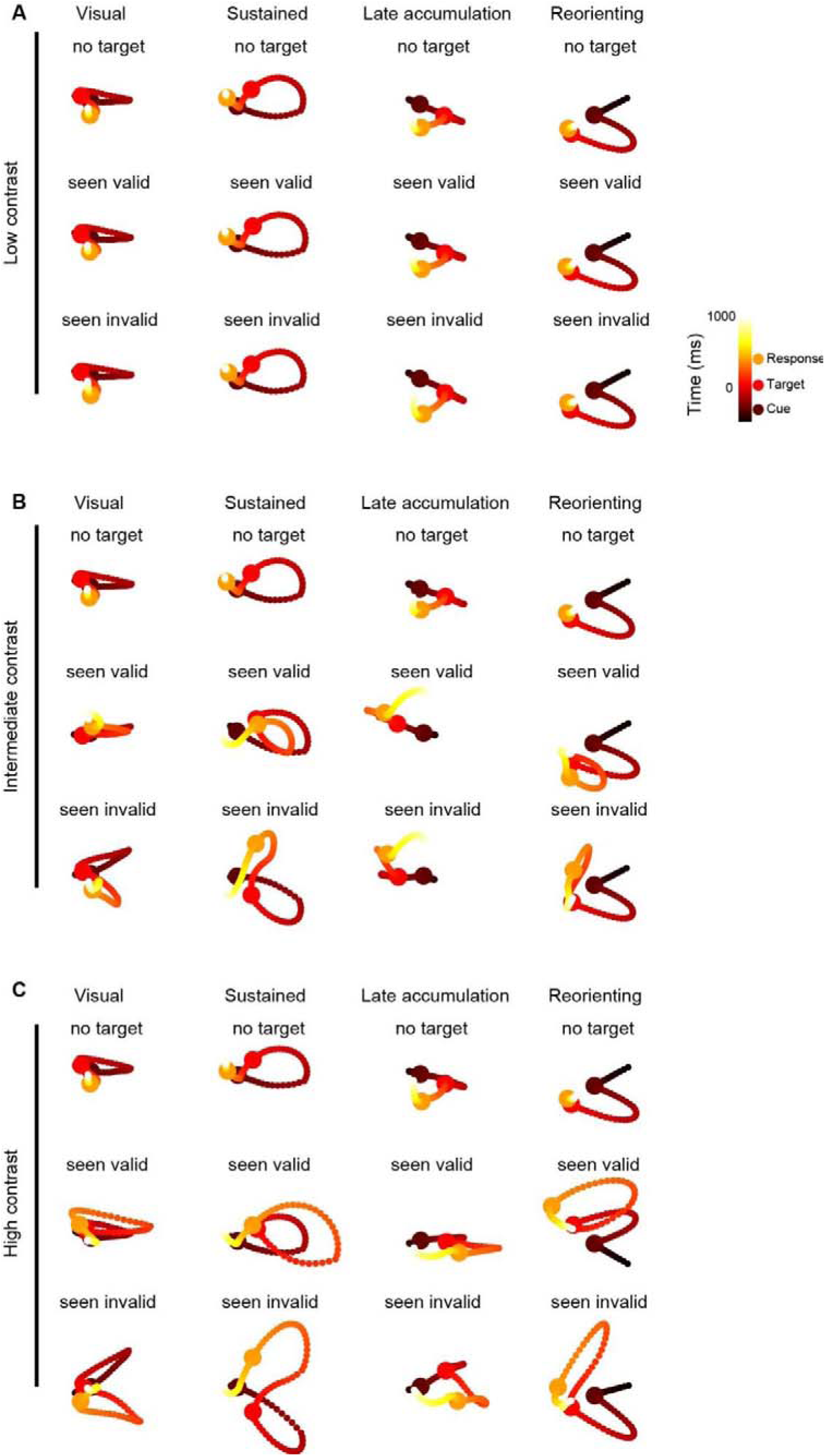
Unit components of RNN clusters. Two-dimensional t-SNE visualization of unit components for (a) low, (b) intermediate, and (c) high target contrast levels by clusters. Color map represents time, darker for earlier and lighter for later time points. Difference in unit component for validly and invalidly cued seen targets emerged when target contrast attained intermediate level, and increased for higher target contrasts.

**Table S1.** Contact localization according to the Desikan-Killiany-Tourville atlas.

| Region name | Implanted<br>Electrode<br>N | Visual<br>N | Sustained<br>N | Late<br>accumulation<br>N | Reorienting<br>N | Conscious<br>report<br>N |
| --- | --- | --- | --- | --- | --- | --- |
| Banks superior temporal sulcus | 4 | 0 | 1 | 0 | 0 | 0 |
| Caudal anterior-cingulate cortex | 11 | 0 | 2 | 0 | 0 | 0 |
| Caudal middle frontal gyrus | 20 | 0 | 2 | 3 | 0 | 2 |
| Cuneus cortex | 1 | 0 | 0 | 0 | 0 | 0 |
| Entorhinal cortex | 5 | 0 | 1 | 0 | 0 | 0 |
| Fusiform gyrus Posterior | 21 | 4 | 2 | 0 | 0 | 6 |
| Fusiform gyrus Middle | 16 | 2 | 3 | 1 | 0 | 2 |
| Fusiform gyrus Anterior | 4 | 0 | 3 | 0 | 0 | 0 |
| Inferior parietal cortex | 39 | 9 | 8 | 1 | 6 | 4 |
| Inferior temporal gyrus Posterior | 16 | 2 | 4 | 2 | 0 | 0 |
| Inferior temporal gyrus Middle | 17 | 0 | 5 | 1 | 1 | 0 |
| Inferior temporal gyrus Anterior | 22 | 0 | 10 | 0 | 0 | 0 |
| Lateral occipital cortex | 12 | 0 | 1 | 0 | 1 | 0 |
| Lingual gyrus | 15 | 0 | 2 | 0 | 0 | 0 |
| Medial orbital frontal cortex | 12 | 0 | 2 | 4 | 0 | 0 |
| Middle temporal gyrus Posterior | 61 | 13 | 15 | 4 | 3 | 2 |
| Middle temporal gyrus Middle | 30 | 0 | 13 | 1 | 1 | 0 |
| Middle temporal gyrus Anterior | 30 | 0 | 2 | 0 | 0 | 0 |
| Parahippocampal gyrus | 7 | 0 | 3 | 3 | 0 | 0 |
| Paracentral lobule | 5 | 0 | 1 | 0 | 0 | 0 |
| Pars opercularis | 10 | 0 | 0 | 1 | 0 | 2 |
| Pars orbitalis | 31 | 1 | 3 | 6 | 2 | 2 |
| Pars triangularis | 8 | 0 | 1 | 3 | 1 | 0 |
| Pericalcarine cortex | 1 | 0 | 0 | 0 | 0 | 0 |
| Postcentral gyrus dorsal | 5 | 0 | 0 | 0 | 0 | 0 |
| Posterior-cingulate cortex | 6 | 2 | 0 | 1 | 0 | 0 |
| Precentral gyrus dorsal | 15 | 0 | 0 | 3 | 0 | 6 |
| Rostral anterior cingulate cortex | 4 | 0 | 1 | 0 | 0 | 0 |
| Rostral middle frontal gyrus | 13 | 0 | 1 | 3 | 0 | 0 |
| Superior frontal gyrus | 73 | 0 | 11 | 18 | 0 | 5 |
| Superior temporal gyrus Posterior | 33 | 1 | 7 | 0 | 3 | 2 |
| Superior temporal gyrus Middle | 22 | 0 | 4 | 0 | 0 | 0 |
| Superior temporal gyrus Anterior | 25 | 0 | 6 | 0 | 0 | 0 |
| Supramarginal gyrus | 22 | 0 | 1 | 4 | 1 | 3 |
| Temporal pole | 19 | 0 | 10 | 0 | 0 | 0 |
| Postcentral gyrus ventral | 6 | 0 | 1 | 0 | 0 | 0 |
| White matter | 62 | 8 | 14 | 5 | 0 | 2 |
| Hippocampus | 24 | 0 | 8 | 3 | 0 | 0 |
| **Total** | **727** | **42** | **148** | **67** | **19** | **38** |
| **Frontal** | 88 | 1 | 24 | 42 | 3 | 18 |
| **Parietal** | 39 | 11 | 10 | 5 | 7 | 6 |

## Notes

### Competing Interest Statement

The authors have declared no competing interest.

### Summary of Updates

revised paper based on received reviewers' comments

